# MitoNEET reduces the mitochondrial oxidative phosphorylation during epithelial-mesenchymal transition

**DOI:** 10.1101/2024.07.29.603210

**Authors:** Haruka Handa, Yasuhito Onodera, Tsukasa Oikawa, Shingo Takada, Koji Ueda, Daiki Setoyama, Takashi Yokota, Miwako Yamasaki, Masahiko Watanabe, Yoshizuki Fumoto, Ari Hashimoto, Soichiro Hata, Masaaki Murakami, Hisataka Sabe

## Abstract

Mitochondrial functions range from catabolic to anabolic, which are tightly coordinated to meet cellular demands for proliferation and motility. MitoNEET is a mitochondrial outer membrane protein with a CDGSH domain and is involved in mitochondrial function. Epithelial-to-mesenchymal transition (EMT) is the process in which cells lose their epithelial characteristics and acquire mesenchymal traits, such as motility, which is a vital step for organism development and wound-healing. Cellular motility is associated with high ATP consumption owing to lamellipodia formation, which is supported by upregulated oxidative phosphorylation (OXPHOS) capacity. However, how mitoNEET is involved in the regulation of OXPHOS capacity and subsequent cellular motility remains unclear. Here we show that loss of mitoNEET regulation during EMT impairs both OXPHOS enhancement and cell motility in non-transformed NMuMG mouse mammary gland epithelial cells. We found that mitoNEET is downregulated during EMT, and that the aberrant expression of mitoNEET abolishes the upregulation of OXPHOS, leading to the inhibition of cell motility. Furthermore, we found that mitoNEET topology may be crucial for the regulation of the mitochondrial electron transfer chain, suggesting an additional regulatory pathway for OXPHOS capacity. Our results demonstrate that mitochondrial OXPHOS capacity during EMT is partly regulated by the dynamics of the outer membrane protein. We believe that our findings are the first step towards understanding the mechanisms by which mitochondrial outer membrane protein topology affects organelle functions.

## Introduction

MitoNEET (mNT/Cisd1) is a dimeric single-span outer mitochondrial membrane (OMM) protein, originally reported as a target of pioglitazone, which is a drug for type- 2 diabetes mellitus (1, 2). MitoNEET function was initially reported to be involved in oxidative capacity by experiments of mitoNEET silencing in mouse heart mitochondria (2). We have recently reported that the age-dependent loss of mitoNEET expression may be a contributing factor to age-associated heart failure, in which OXPHOS capacity is impaired (3). However, the mechanisms by which the expression of mitoNEET is downregulated during the ageing process remain unclear. MitoNEET has a CDGSH domain in its C-terminus, protruding towards the cytosol, which has been reported to function to hold and transfer 2Fe-2S clusters to acceptor proteins (2, 4, 5). Other research has demonstrated that mitoNEET utilises 2Fe-2S clusters to sense redox conditions for the gating of voltage-dependent anion channel 1 (VDAC1), which functions as a system of cytosol-mitochondria communication (6). Regarding mitochondrial dynamics, the predisposition of mitoNEET to dimerization could be a key factor in the development of intermitochondrial junctions, which are essential for maintaining mitochondrial integrity (7). Numerous studies have supported that the CDGSH domain of mitoNEET is important for mitochondorial function and homeostasis. However, the discrepancy between the location of CDGSH domain of mitoNEET (cytosol) and the site of OXPHOS (inner mitochondrial membrane: IMM) evokes some question about how mitoNEET directly affects OXPHOS. Therefore, this question might become a rationale for investigating the possibility that not all but a population of mitoNEET directly associates with the OXPHOS machinery, with its domain facing towards the inter membrane space (IMS).

As eukaryotic cells undergo transdifferentiation into other cell types, metabolic shifts play a pivotal role in maintaining cell integrity and functionality. Epithelial-to-mesenchymal transition (EMT) is a dynamic process in which epithelial cells lose their epithelial characteristics and acquire mesenchymal characteristics, which is represented by motility. This is driven by EMT transcription factors (TFs) such as Zeb1, Snail and Slug (8). It has been well documented that these TFs confer EMT-induced cell motility, involving cytoskeletal remodelling and the expression of extracellular adhesion molecules. Metabolomic analyses, mostly in cancer EMT cells, have shown that these mesenchymal cells use mitochondrial metabolic machinery for their invasive properties and high proliferation rate by altering the expression levels of metabolic enzymes (9). However, other mechanisms by which mitochondrial proteins other than metabolic enzymes regulate their function during EMT are still unclear.

To gain a better understanding of mitoNEET, we first confirmed the downregulation of mitoNEET during EMT and then analysed OXPHOS capacity using the Oxygraph-2k. Our aim is to elucidate the association between EMT and mitoNEET and to understand how mitoNEET affects mitochondrial function. To this end, we have performed immunoprecipitation followed by mass spectrometry (IP-MS) analysis to identify proteins interacting with mitoNEET. Our findings suggest that mitoNEET faces not only the cytosol, but a propotion of the protein also faces the IMS. The results of our study will demonstrate that the topology of mitoNEET is a crucial factor in elucidating its effects on OXPHOS capacity during EMT.

## Materials and methods

### Cell culture

NMuMG cells, 3T3-Swiss and A549 were purchased from American Type Culture Collection. NMuMG cells were cultured in high glucose Dulbecco’s Modified Eagle Medium (DMEM) (Nacalai Tesque) with 10% Fetal Bovine Serum (FBS) (Biowest) and 1/1000 of 10 mg/mL Insulin (FUJIFILM Wako Pure Chemical Corporation). A549 and 3T3-Swiss cells were cultured in high glucose DMEM with 10% FBS, and 5 µg/mL of human Transforming growth factor (TGF) -β (Peprotech) was added to the culture medium for the induction of epithelial-mesenchymal transition (EMT).

### siRNA and transfection

The siRNAs targeting mitoNEET were purchased from horizon (ON-TARGETplus, SMARTPool). Cells were transfected with siRNA duplexes using DharmaFECT 1 (horizon), according to the instruction provided by the manufacturer. Assays were performed 48 hours post-transfection, otherwise stated.

### Wound-healing assay

A total of 1.0 × 10^4^ of NMuMG cells were seeded on to each well of the ibidi Culture-Insert 2 Well (ibidi) and induced with or without TGF-β the next day. Two days after stimulation, cells were stained with Hoechst33342 (Dojindo) for 30 minutes and the insert was removed to start wound-healing. The wound closure process was recorded using Nikon A1R for 24 hours. Data were analysed with NIS-elements software (Nikon).

### Transwell assay

A total of 3.0 × 10^4^ of NMuMG cells induced with or without TGF-β for two days were seeded on the upper wells of an 8-µm pore size transmembrane (Falcon #353097), and incubated in the presence or absence of 10% FBS for 4 hours. A total of 8.0× 10^4^ Swiss- 3T3 cells were seeded onto the upper wells 2 days post siRNA transfection and incubated for 6 hours. Membranes were fixed in 4% paraformaldehyde in phosphate buffered saline (PBS) (Nacalai Tesque), and soaked in 0.4% crystal violet solution. Images of six microscopic fields of the region covered with stained cells that migrated out to the lower surface were obtained by using EVOS® FL Cell Imaging System (ThermoFisher Scientific). Then the area was measured by Image J software as ‘% migration area’.

### Antibodies and immunoblotting analyses

The following antibodies were purchased from commercial sources: mitoNEET (Cell Signaling Technology [CST] #83775 or Proteintech #16006-1-AP), E-cadherin (BD Transduction Laboratories #610182), N-cadherin and β-actin (EMD Millipore), Smad2 (CST #3103), Ser465/467-phosphorylated Smad2 (CST #3108), Tomm70 (Proteintech #14528-1-AP), Timm50 (Proteintech #22229-1-AP), and Prohibitin (Proteintech #10787-1-AP). Electrophoresed proteins were transferred onto polyvinylidene difluoride membranes, as described previously (10). β-actin was used as a loading control. NeutrAvidin 800 (Invitrogen #SA535521) was used for the detection of biotin. ECL kit (GE Healthcare) and SuperSignal West Dura (Thermo Fisher Scientific) were used to detect specific proteins, whereas Odyssey (LI-COR) was used to simultaneously detect a specific protein and biotin.

### Cell lysis for Protein extraction

Cells were lysed by RIPA buffer (150 mM NaCl, 20 mM Tris-HCl [pH 7.4], 5 mM ethylenediaminetetraacetic acid (EDTA), 1% NP-40, 1% sodium deoxycholate, and 0.1% sodium disulfide (SDS)) added with Halt™ protease inhibitor cocktail (ThermoFisher Scientific) and incubated on ice for 10 min, followed by centrifugation at 17,900 *g* and 4°C for 30 min. The supernatant was used as the whole cell lysate.

### Reverse transcription (RT) and quantitative RT-polymerase chain reaction (qRT-PCR)

Total RNA was extracted from cultured cells using RNeasy Plus Mini kit (QIAGEN), according to the manufacture’s instruction. RNAs were reverse-transcribed by SuperScript VILO Master Mix (Invitrogen), and 50 ng of RNAs were used for qRT-PCR. qRT-PCR reactions were performed in duplicate using TaqMan Universal PCR Master Mix and TaqMan gene expression assays (#4331182, Applied Biosystems), and measured by the 7300 Real Time PCR System (Applied Biosystems). The following TaqMan probes (Applied Biosystems) were used: mouse *Cisd1* Mm01172641_g1; human *CISD1* Hs05024086_m1; mouse *Gapd* #4351309; mouse *Actn* Mm02619580_g1; human *GAPDH* Hs02758991_g1; human *ACTN* Hs01060665_g1. Relative quantity values (RQ) of the samples were calculated by the delta Ct method. Then, geometric means of RQs (GeoRQ-ref) of two reference genes, *Gapdh* and *Actb*, were used to normalise the genes of interest. Finally, the following equation was used to calculated relative gene expressions: RQ-GOI ÷ GeoRQ-ref.

### Primers, vectors and transfection

PCR was performed to amplify cDNAs encoding full-length mouse mitoNEET from NMuMG mRNAs extracted by RNeasy Plus Mini kit (QIAGEN). HA-tag was attached in the C-terminus of mitoNEET with the insertion of two glycine amino acids. *mitoNEET-HA* was then inserted after the promoter of pPB-CEH-MCS-IP vector (11). All primers used for subcloning are listed in the table 1. An irrelevant sequence (AGCGAGGTTTACATGTTGTGTGATAGTGAAGCCACAGATGTATCACACAACATGTAAACCA) was also inserted into the same location as a control. MicroID2 (12) with a sequence which is designed to target mitochondrial intermembrane space (IMS) was inserted in doxycycline-inducible pPB vector encoding blasticidin-resistance protein. For stable transfection, cells were co-transfected with pPB vectors and a plasmid encoding hyperactive piggyBac transposase by Viafect (Promega) as described previously (11). Cells were selected by puromycin (2 µg/mL) and/or blasticidin (5 µg/mL) for 7 days.

**Table 1.**
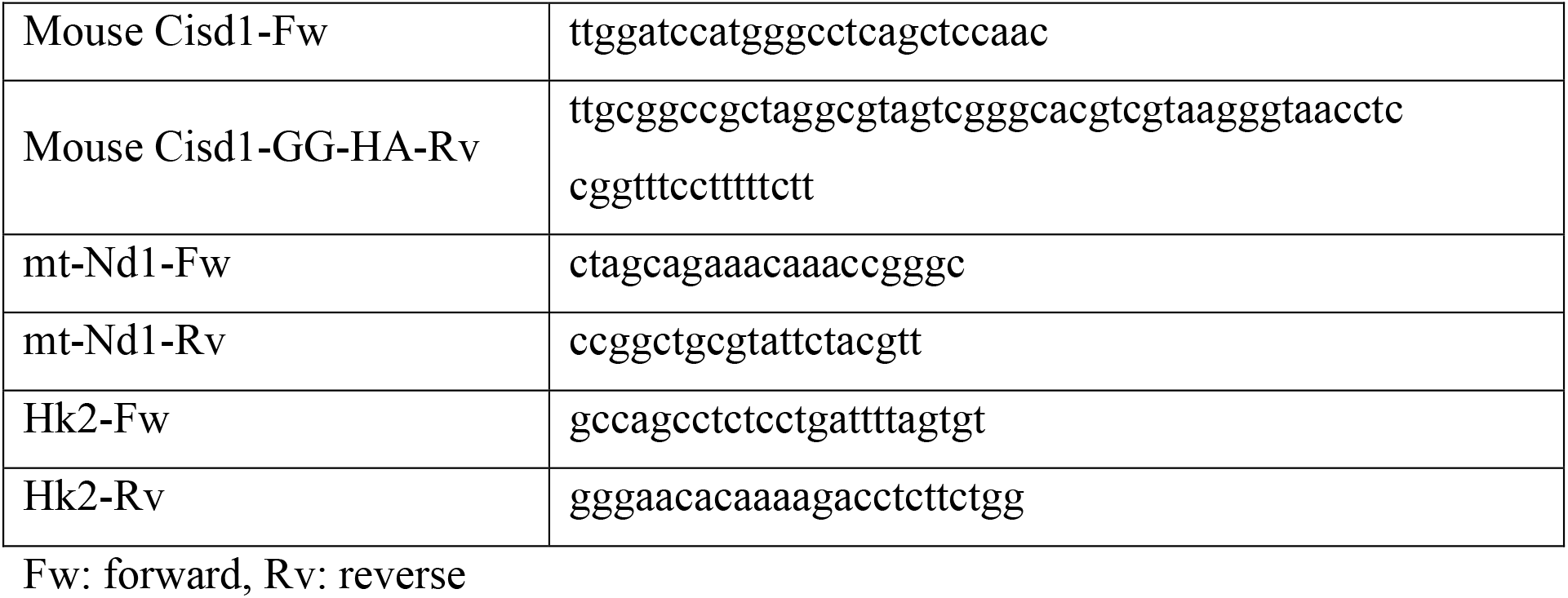
(Primers)

### Biotinylation assay

MicroID2 was induced with 100 ng/mL Doxycycline (FUJIFILM Wako Pure Chemical Corporation) for two days and 50 µM D-biotin (Nacalai Tesque) was added to the culture medium 6 hours prior to cell lysis.

### Immunoprecipitation

After quickly washed with PBS on ice, cells were lysised with immunoprecipitation (IP) solution (10 mM Tris-Cl [pH 7.4], 100 mM NaCl, 1% NP-40 (Nacali Tesque), 1% digitonin (Nacalai Tesque), and 5% Glycerol with Halt™ protease inhibitor cocktail (ThermoFisher Scientific) at 4°C for 15 min. After clarifying by centrifugation, 1,000 µg of cell lysate was incubated at 4°C overnight with 30 µg of Protein G magnetic beads (Tamagawa Seiki) covalently bound to HA-antibody (Sigma) using BS3, a cross- linker (Dojindo). After washing with IP-wash buffer (10 mM Tris-Cl (pH 7.4), 100 mM NaCl, 0.02% Tween-20, and 5% Glycerol) three times, beads were resuspended in Laemmli buffer and boiled for 5 min, and eluted proteins were analysed by SDS-poly acrylamide gel electrophoresis (PAGE).

### Mass spectrometric (MS) analysis of proteins

After reduction with 10 mM TCEP at 100°C for 10 min, and alkylation with 50 mM iodoacetamide at ambient temperature for 45 min, protein samples were subjected to digestion with Trypsin/Lys-C Mix (Promega) at 47°C for 2 hours on S-Trap columns (ProtiFi). The resulting peptides were extracted from gel fragments and analyzed using Orbitrap Fusion Lumos mass spectrometer (Thermo Scientific) combined with UltiMate 3000 RSLC nano-flow HPLC (Thermo Scientific). Peptides were enriched using μ-Precolumns (0.3 mm i.d. × 5 mm, 5 μm; Thermo Scientific) and separated on AURORA columns (0.075 mm i.d. × 250 mm, 1.6 μm, Ion Opticks Pty Ltd.) using the following two-step gradient: 2% to 40% acetonitrile for 110 min, followed by 40% to 95% acetonitrile for 5 min in the presence of 0.1% formic acid. The compensation voltages for gas-phase fractionation using FAIMS Pro (Thermo Scientific) were set at −40, −60, and −80 V. The analytical parameters of Orbitrap Fusion Lumos were set as follows: Resolution of full scans = 50,000, Scan range (m/z) = 350 to 1,500, Maximum injection time of full scans = 50 msec, AGC target of full scans = 4 × 10^5^, Dynamic exclusion duration = 30 sec, Cycle time of data dependent MS/MS acquisition = 2 sec, Activation type = HCD, Detector of MS/MS = Ion trap, Maximum injection time of MS/MS = 35 msec, and AGC target of MS/MS = 1 × 10^4^.

The MS/MS spectra were searched against the *Mus musculus* protein sequence database in SwissProt using Proteome Discoverer 3.0 software (Thermo Scientific), in which peptide identification filters were set at “false discovery rate < 1%”. Label-free relative quantification analysis for proteins was performed with the default parameters of Minora Feature Detector node, Feature Mapper node, and Precursor Ions Quantifier node in Proteome Discoverer 3.0 software.

### Mitochondria isolation

Mitochondria were roughly isolated as previously described with some modifications (13). Cells were swelled by incubating in hypotonic buffer (10 mM Tris-MOPS [pH7.4], 1 mM EDTA-Tris [pH7.4]) with rotation at 4°C for 30 min. The swelled cells were homogenised by 20 repeated passes through a 28-gauge needle attached to a 1-mL syringe. The homogenate was centrifuged at 700 *g* and 4°C for 10 min, and the supernatant was centrifuged again at the same condition, followed by another centrifugation at 7,000 *g* and 4°C for 10 min. The pellet was suspended in isotonic buffer (10 mM Tris-MOPS [pH7.4], 1 mM EDTA-Tris [pH7.4], and 200 mM sucrose) and was centrifuged at 7,000 *g* and 4°C for 10 min. The pellet was again suspended in isotonic buffer and was centrifuged at 10,000 *g* and 4°C for 10 min. The pellet was then resuspended in RIPA buffer and incubated on ice for 10 min. The lysate was centrifuged at 17,900 *g* and 4°C for 30 min. This supernatant was used as the mitochondrial fraction.

### Mitochondrial oxygen consumption rate (OCR) analysis

The mitochondrial respiratory capacity was measured at 37°C using a high-resolution respirometer (Oxygraph-2k, Oroboros Instruments). A total of 2.0 × 10^6^ of NMuMG cells were added to the respirometer chamber filled with 2 mL of MiR05 medium (110 mM sucrose, 60 mM potassium lactobionate, 0.5 mM ethylene glycol tetraacetic acid, 3 mM MgCl2, 20 mM taurine, 10 mM KH2PO4, 20 mM HEPES [pH 7.1], and 1% BSA).

The cells were permeabilized with 5 µM digitonin (Merck). Then, substrates of OXPHOS and ADP were added to the respirometer chamber in the following order: (1) 2 mM malate, 10 mM glutamate, and 5 mM pyruvate (complex I-linked substrates); and (2) 5 mM ADP + 3 mM MgCl2; and (3) 10 mM succinate (complex II-linked substrates). Cytochrome *c* was added after (2) to check mitochondrial integrity. Samples with a 10% increase in OCR post cytochrome *c* addition were considered as containing damaged mitochondria, and were discarded. The OCR was expressed as O2 flux normalised to 1 × 10^6^ cells. Data acquisition and analysis of data were performed using DatLab software (Oroboros Instruments), as described previously (14).

### Quantative PCR of mitochondrial DNA and nuclear DNA

Mitochondrial DNA and nuclear DNA and were extracted using NucleoSpin® DNA RapidLyse (Takara). Mouse *NADH dehydrogenase* (*mt-Nd1*) was used as a mitochondrial marker and mouse *hexokinase 2* (*Hk2*) as a nuclear marker. qPCR reactions were performed in duplicate, using primers for *mt-Nd1* and *Hk2* and THUNDERBIRD qPCR Mix (TOYOBO), and measured using the 7300 Real Time PCR System (Applied Biosystems). Primers for *mt-Nd1* and *Hk2* are listed in Table 1.

### Native-PAGE and In-gel activity assay

Isolated mitochondria were incubated with 5% digitonin and NativePAGE^TM^ Sample buffer (ThermoFisher Scientific), followed by electrophoresis on a Clear-Native PAGE gel (ATTO). Total samples were stained using Coomassie brilliant blue R (Sigma). The gels were incubated with substrates for complex I, II, and IV and colouring agent (15).

### Trypsin assay

Isolated mitochondria were gently mixed with trypsin (Gibco) to a final concentration of 0.025% or 0.05%, with or without 1% Triton-X100 (Sigma), followed by incubation at 37°C. The ratio of treatment strength was calculated by using the concentration of trypsin and the duration of incubation.

### Extraction and liquid chromatography–mass spectrometry (LC-MS) analysis of tricarboxylic acid (TCA)-cycle intermediates and water-soluble metabolites

TCA cycle intermediates were extracted from whole cell pellets by the modified Bligh and Dyer procedure. Briefly, cell pellets were added to 500 µL of ice-cold 80% methanol, vortexed, sonicated five times (30-s sonication and cooling) using BIORUPTOR II (CosmoBio), and centrifuged at 21,500 × *g* for 5 min at 4°C. Supernatants were collected, added with 0.8 volume of chloroform and 0.5 mL water were added, vortexed, and centrifuged at 21,500 × *g* for 5 min at 4°C. the upper water layer was dried using an miVac DUO concentrator system (SCRUM Inc.). The resulting metabolites were dissolved in 100 µL of solvent A and analysed using a triple quadrupole mass spectrometer (LCMS-8040, Shimadzu). To monitor TCA-cycle metabolites, reverse phase ion-pair chromatography was performed using an ACQUITY UPLC BEH C18 column (100 × 2.1 mm, 1.7 µm particle size; Waters). The mobile phase consisted of solvent A (15 mM acetic acid and 10 mM tributylamine in 3% methanol) and solvent B (methanol), and a column oven temperature of 40℃. The gradient elution program was as follows: a flow rate of 0.3 mL/min: 0–3 min, 0% B; 3– 5 min, 0%–40% B; 5–7 min, 40%–100% B; 7–10 min, 100% B; and 10.1–14 min, 0% B. Parameters for the negative electrospray ionization source (ESI) mode under multiple reaction monitoring (MRM) were as follows: drying gas flow rate, 15 L/min; nebulizer gas flow rate, 3 L/min; DL temperature, 250℃; and heat block temperature, 400℃; and collision energy (CE), 230 kPa. To measure wide varieties of water-soluble metabolites, the supernatants described above were diluted 10-fold with 0.1% formic acid and separated on a Discovery HS-F5-3 column (150 × 2.1 mm, 3 μm particle size; Sigma-Aldrich) with mobile phases consisting of solvent A (0.1% formic acid) and solvent B (0.1% formic acid in acetonitrile). The column oven temperature was 40℃. The gradient elution program was as follows: a flow rate of 0.25 mL/min: 0–2 min; 0%B; 2– 5 min, 0%–25%B; 5–11 min, 25%–35%B; 11–15 min, 35–95%B; 15–25 min, 95%B; and 25.1–30 min, 0%B. The parameters for the heated electrospray ionization source (ESI) in negative/positive ion mode under multiple reaction monitoring (MRM) were as follows; drying gas flow rate, 10 L/min; nebulizer gas flow rate, 3 L/min; heating gas flow rate, 10 L/min; interface temperature, 300℃; DL temperature, 250℃; heat block temperature, 400℃; and CID gas, 270 kPa.

### Extraction and LC-MS analysis of acylcarnitine

Whole cell pellets were mixed with ice-cold 80% methanol, vortexed, sonicated five times (30-s sonication and 30-s cooling) with BIORUPTOR, and centrifuged at 21,500 *g* for 5 min at 4℃. The supernatants were analysed using a triple quadrupole mass spectrometer (LCMS-8060, Shimadzu). To monitor acylcarnitines, HILIC chromatography was performed using a Luna 3u HILIC 200A column (150 × 2.0 mm, 3 µm particle size, Phenomenex). The mobile phase consisted of solvent A (10 mM ammonium formate) and solvent B (10 mM acetonitrile: 10 mM ammonium formate, 9:1), and the column oven temperature was 40℃. The gradient elution program was as follows: a flow rate of 0.3 mL/min: 0–2.5 min, 100% B; 2.5–4 min, 100%–50% B; 4–7.5 min, 50%–5% B; 7.5–10 min, 5% B; and 10.1–12.5 min, 100% B. The parameters for the heated ESI in the positive ion mode under precursor ion scan were as follows: drying gas flow rate, 10 L/min; nebulizer gas flow rate, 3 L/min; heating gas flow rate, 10 L/min; interface temperature, 300°C; DL temperature, 250°C; heat block temperature, 400°C; and CID gas, 270 kPa.

### Data processing and metabolomics data analysis

Data processing was carried out using the LabSolutions LC-MS software (Shimadzu, Japan). Then processed data were analysed by MetaboAnalyst (16).

### Immunofluorescence microscopy

Cells were seeded onto µ-slides (ibidi) and fixed in 4% paraformaldehyde in PBS (Nacalai Tesque). To observe mitochondria, cells were transfected with pPB-mitoGFP2 vector, and incubated for 2 days before fixation. The fixed cells were permeabilised with 0.1% Triton X-100 in PBS for 5 min, washed with PBS, and then incubated with Maxblock Blocking Medium (Active Motif) at room temperature for 2 hours. Samples were then subjected to immunostaining using anti-HA antibody, followed by a Cy3- conjugated secondary antibody. The samples were sealed with ProLong™ Glass Antifade Mountant with NucBlue™ Stain (ThermoFisher). Fluorescence images were acquired using a confocal laser-scanning microscope (Model A1R with NIS-Elements software, Nikon) using a Plan-Apo VC 100× lens (Nikon) and analysed with the attached software.

### Postembedding immunogold electron microscopy

After removing the medium from culture dishes, fixative (4% paraformaldehyde and 0.1% glutaraldehyde in 0.1 M phosphate buffer (PB) [pH 7.2]) was quickly applied and left *in situ* for 15 min at room temperature. Cells were scraped, transferred to a 1.5 ml Eppendorf tube, and postfixed in the same fixative for 2 hours. Samples were centrifuged at 11,750 *g* for 3 min at 4°C, and the supernatant was removed. Pellets were washed with 0.1 M PB, cryoprotected by immersion in 30% glycerol in 0.1M PB for 30 min, and frozen rapidly in liquid propane (–180°C) in an EM CPC unit (Leica Microsystems). Frozen pellets were immersed in 0.5% uranyl acetate in methanol at −90°C in an AFS freeze-substitution unit (Leica Microsystems), infiltrated at −45°C with Lowicryl HM-20 resin (Electron Microscopy Sciences) and polymerized with UV for 48 hours. Ultrathin sections (90-nm thick) were cut using a Leica EM UC7 ultramicrotome (Leica Microsystems), and mounted onto nickel grids.

Ultrathin sections on nickel grids were etched with saturated sodium-ethanolate solution for 1–5 s, incubated in tris buffered saline (TBS) containing 0.03% Triton X-100 (TBS- T), 0.1% sodium borohydride, and 50 mM glycine for 10 min, and treated with the following successive solutions: the goat blocking solution containing 2% normal goat serum (Nichirei) in TBST for 20 min, and rabbit anti-HA mAb (#3724, Cell Signaling Technology; 10 μg/mL diluted with the blocking solution) overnight. After washing with TBS-T three times, grids were incubated with colloidal gold (5 nm in diameter)- conjugated anti-rabbit IgG (EM.GAR5, British BioCell International; 1:100 in the blocking solution) for 2 hours. Finally, grids were washed with TBS-T and distilled water, and stained with 1% OsO4 for 15 min, 5% uranyl acetate/40% ethanol for 90 s, and Reynold’s lead citrate solution for 1 min. Photographs were taken with a JEM1400 electron microscope (JEOL) at the original magnification 50,000× For quantitative analysis, immunogold particles that fall within 30 nm of the OMM or IMM were selected, and the distance between the OMM was measured using MetaMorph software (Molecular Devices).

### Statistical Analysis

All data except for metabolome and proteome data were analysed using Prism 10 (GraphPad). Data are expressed as the mean ± error bars. Error bars indicate standard error of the mean (SEM). Points indicate each experimental datum. Welch’s *t*-test was used for comparison of two samples and multiple comparison test with Tukey’s or Dunnett’s correction if the experiment had more than two samples unless otherwise stated. *P*-values and *adjusted P*-values (*adj. P*) less than 0.05 were considered to indicate statistically significant differences between groups.

## Results

### Some population of mitoNEET faces towards intermembrane space of mitochondrion

We previously reported that mitoNEET might function as a positive regulator of OXPHOS capacity in cardiac myocyte (3). However, it is still unclear how mitoNEET can regulate OXPHOS capacity. mitoNEET has a transmembrane domain to insert itself into the outer mitochondrial membrane (OMM) (Fig.1A). On the other hand, inner mitochondrial membrane (IMM) is a base for ETC and OXPHOS proteins (17). Therefore, if mitoNEET regulates the OXPHOS capacity, it is reasonable to conjecture that CDGSH domain is facing toward the mitochondrial intermembrane space (IMS). To examine this possibility, we generated the mitoNEET overexpression cell line, NMuMG mNT-HA cells, which stably expresses mitoNEET tagged with HA in its C-terminus, and performed co-immunoprecipitation (IP) followed by mass spectrometry (MS). As a control, we used irrelevant sequence expressing NMuMG cells (Irr). Some proteins known as an associating partner of mitoNEET, such as Ciao1, were detected (Fig. 1B). On the other hand, we found that some of the listed proteins are reported to localise in another place including IMM (Fig. 1B). We therefore hypothesised that mitoNEET might have its CDGSH domain facing not only the cytoplasm but also the IMS, allowing it to interact with those IMM proteins. To confirm the topology of mitoNEET, we performed the trypsin digestion assay against wild-type NMuMG cells, in which trypsin digests the proteins on the outer side of OMM including mitoNEET. As expected, Tomm70, known as an OMM protein, was digested according to the concentration of trypsin, whereas Timm50, an IMM protein, was detected at almost the same level as in the non-digested mitochondria (Fig. 1C). In this assay, we used an anti- mitoNEET antibody whose targeting epitope was around Asp61 of mitoNEET and was reported to face towards cytosol. mitoNEET has amino acid residues that can be cleaved with trypsin if facing to the cytosol. Thus, it should be noted that if mitoNEET faces towards cytosol, it is not detected with this antibody after digested by trypsin. This result insisted that at least a portion of mitoNEET is thought to be face towards IMS (Fig. 1C). We next developed another method to assess mitoNEET topology using MicroID2 expressed in the IMS. Both NMuMG cell lines (Irr and mNT-HA) were transfected with the doxycycline-inducible MicroID2 vector. When precipitated by neutravidin beads, both Timm50 and mitoNEET were detected in the biotinylated proteins. (Fig. 1D). This strongly supports that some population of mitoNEET is facing towards the IMS. To exclude the possibility that mitoNEET exists in the IMM, we utilized immunogold electron microscopy to analyse the distribution of mitoNEET localization within mitochondria (Fig. 1E). For this purpose, we used an anti-HA antibody to label mitoNEET. Considering that the size of an IgG antibody is 15 nm and a gold colloid is 5 nm, the maximum positional error for each immunogold-labelled molecule is estimated to be 20 nm. Under these conditions, immunogold labelling for mNT-HA demonstrated a broad but distinct distribution (Fig. 1E, upper). When we plotted the fraction of immunogold particles against the distance from the OMM (Fig. 1E, lower), we observed that the distribution for mNT-HA was concentrated on the OMM. These results show that most mitoNEET molecules exist on the OMM and not on the IMM, supporting our hypothesis that a population of mitoNEET faces towards the IMS to regulate OXPHOS capacity.

**Figure 1.**
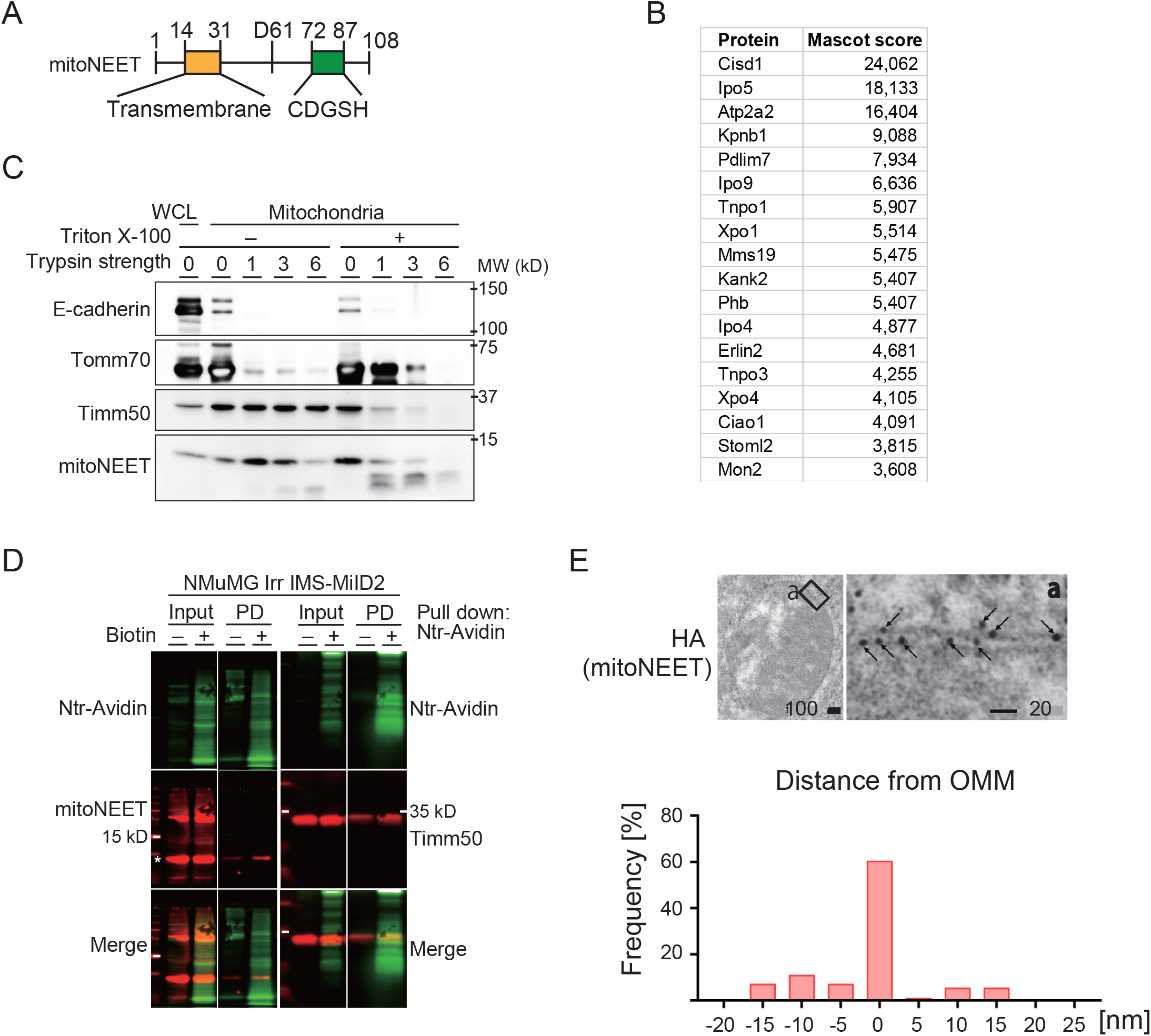
A population of mitoNEET faces the intermembrane space of mitochondria. A, Schematic illustration of the mitoNEET protein. The transmembrane domain and CDGSH domain are depicted. **B,** List of top 18 proteins detected by IP-MS using the HA-antibody in NMuMG mNT-HA cells. **C,** Immunoblots of the indicated antibodies in trypsin-digested crude mitochondria with or without Triton X-100. WCL: whole cell lysates. **D,** SDS-PAGE followed by immunoblotting (red) and NeutrAvidin blotting (green) of the indicated proteins. Lysates from NMuMG cells transfected with a vector coding for MicroID2 that is targeted to the inter mitochondrial space (IMS) (IMS-MiID2), with or without biotinylation, were pulled down by NeutrAvidin beads (Ntr-Avidin). PD: pull down. **E,** Representative images of mitochondria stained using the indicated antibodies (upper) and a graph showing the distribution of 5-nm Au beads across the mitochondrial membrane (lower). The unit of the scale bars is nm. IMM: inner mitochondrial membrane, OMM: outer mitochondrial membrane. The calculated intermembrane distance was 9.4 ± 1.7 nm.

## OXPHOS capacity is enhanced during EMT in NMuMG cells

As mitoNEET has been demonstrated to have a positive effect on OXPHOS capacity in cardiomyocytes (Furihata, 2021), we also confirmed whether mitoNEET regulates OXPHOS capacity in epithelial cells and fibroblasts. The capacity of OXPHOS was measured according to the substrate-uncoupler-inhibitor-titration (SUIT) protocol, in which complex I-linked and complex I + II-linked substrates were added (Sup. 1A). Unexpectedly, *mitoNEET* silencing by siRNA did not affect OXPHOS capacity in two types of cell lines, namely, NMuMG cells and Swiss 3T3 cells, which have epithelial and mesenchymal characteristics, respectively (Fig. 2A and 2B). These results indicate that mitoNEET has negligible effects on OXPHOS capacity in epithelial cells and fibroblasts, at least in normal culture conditions. On the other hand, *mitoNEET* has been reported to be downregulated during epithelial-mesenchymal transition (EMT) (public dataset: GSE103372). We confirmed that mitoNEET is downregulated at both the protein and mRNA levels by inducing EMT in NMuMG cells (Fig. 2C and 2D). This downregulation was also found in A549 cells that were induced to undergo EMT (Sup. 1B, 1C, and 1D). EMT was morphologically confirmed in those cells at 48 hours post-stimulation (Fig. 2E). We found that in NMuMG cells, OXPHOS capacity was enhanced 48 hours post TGF-β stimulation (Fig. 2F). Therefore, we hypothesised that mitoNEET negatively regulates the OXPHOS capacity during EMT.

**Figure 2.**
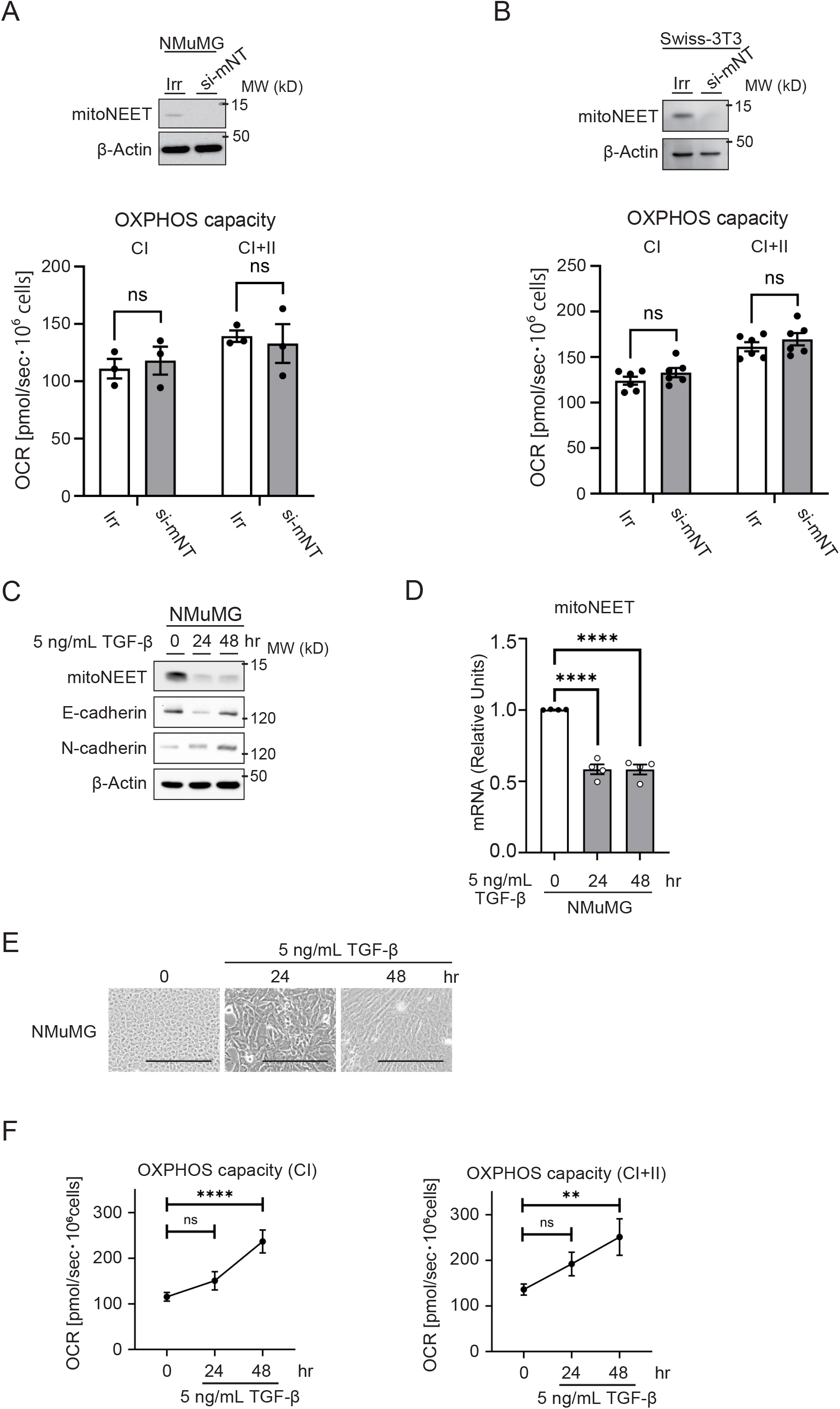
OXPHOS capacity is enhanced during EMT in NMuMG cells. A, Immunoblots of mNT (top), and OXPHOS capacity in the presence of Complex I (CI)- or CI + II-linked substrates (bottom) of siRNA-treated NMuMG cells. **B,** Immunoblots of mNT (top), and OXPHOS capacity of siRNA treated Swiss-3T3 cells. (In A and B, ns means statistically nonsignificant by Welch’s *t*-test.) **C,** Immunoblots of NMuMG cells pre and post TGF-β stimulation. **D,** Levels of mitoNEET mRNA in TGF-β-stimulated NMuMG cells measured by RT-qPCR. **E,** Representative images of morphological changes in NMuMG cells stimulated by 5 ng/mL TGF-β. Bars, 200 µm. **F,** Chronological alterations post TGF-β stimulation in OXPHOS capacity of mitochondria from NMuMG cells. (D and F: Dunnett’s multiple comparison test, \*\*\*\**adj. P* < 0.0001, ** *adj. P* < 0.005, * *adj. P* < 0.05.)

## MitoNEET suppresses the enhancement of OXPHOS capacity and cell motility during EMT

To investigate the possibility that mitoNEET acts as a negative regulator during EMT, we used mNT-HA NMuMG cells and Irr cells that were induced to undergo EMT. We also confirmed that overexpressed mitoNEET is localised in mitochondria, as was previously reported (Fig. 3A) (2). The overexpression of mitoNEET did not affect the morphological transformation or the expression of EMT markers, such as phosphorylated Smad2 and E-cadherin induced by TGF-β stimulation (Fig. 3B and 3C). We found that the enhancement of OXPHOS capacity during EMT was cancelled in mNT-HA cells constitutively expressing mitoNEET (Fig. 3D, left). This cancelation was correlated with the lower level of ATP generation in mNT-HA cells than in Irr cells (Fig. 3D, right). Metabolome analysis of these cells also indicated the enhancement of OXPHOS in Irr cells and its cancelation in mNT-HA cells, since TCA metabolites were detected at higher levels in Irr cells than in mNT-HA cells (Sup. 1E). Therefore, the continued expression of mitoNEET upon EMT might suppress the OXPHOS capacity. In addition, we confirmed that the ratio of mitochondria DNA and nuclear DNA was not significantly altered (Fig. 3E), and the amounts of complexes I, II, and IV were not apparently changed, as analysed by the in-gel assay (Sup. 1F). Thus, these results indicate that the cancelation of enhanced OXPHOS capacity during EMT in the presence of mitoNEET is not owing to alterations in mitochondrial quantity. We then assumed that the continuously expressed mitoNEET proteins can interact with the ETC and OXPHOS-associated proteins. To assess this possible interaction during EMT, we focused on the inner membrane protein, prohibitin (Phb), which was listed in the IP-MS data (Fig.1), and was reported to contribute to ETC activity (18). In this experiment, we also used the anti-mitoNEET antibody for precipitation of the endogenous mitoNEET protein, in addition to the anti-HA antibody for precipitation of overexpressed mNT-HA. Phb was precipitated in both Irr and mNT-HA cells stimulated with TGF-β (Fig. 3F). These data also bolster our findings that some mitoNEET faces the IMM, and not the cytosol. Moreover, this interaction between mitoNEET and IMM proteins may explain the inhibitory effect of mitoNEET against the OXPHOS enhancement during EMT, in which the detailed mechanism remains unclear at present. We next investigated the biological outcome of the OXPHOS enhancement. It has been demonstrated that cell motility requires energy production in the form of ATP for the rearrangement of cytoskeleton components, such as actin stress fibres (19). From this viewpoint, we assessed the motility of Irr and mNT-HA cells, pre and post TGF-β stimulation. As we expected, mNT-HA cells had lower motility in wound healing and 3D-chemotaxis assays than Irr cells (Fig. 3G), even though EMT was induced in both cell lines (Fig. 3B and 3C). These data support that mitoNEET needs to be downregulated for the EMT-induced enhancement of OXPHOS capacity.

**Figure 3.**
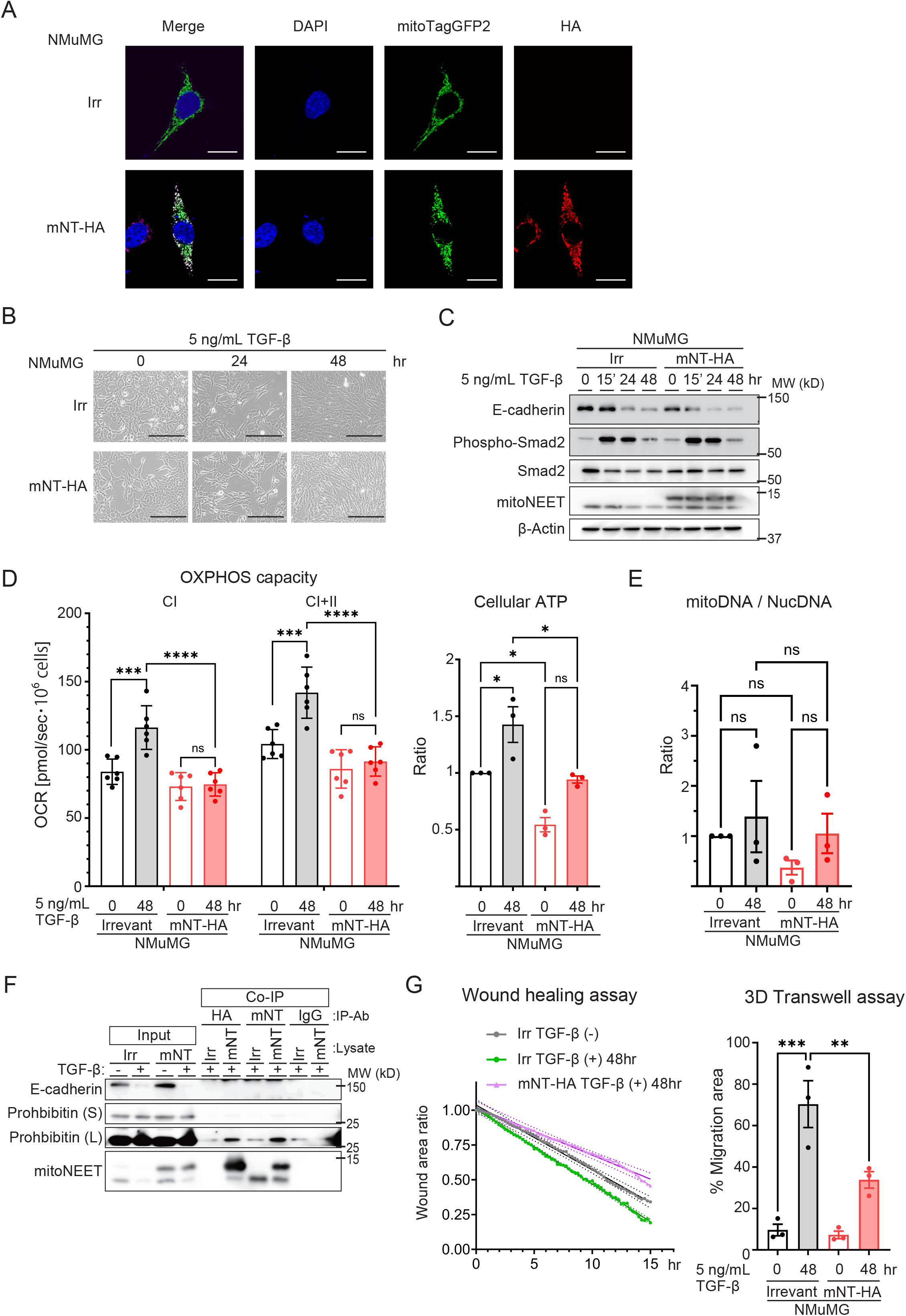
MitoNEET suppresses the enhancement of OXPHOS capacity and cell motility during EMT. A, Representative images of NMuMG Irr and mNT-HA localisation in NMuMG cells. Mitochondria were observed using transiently expressed mitoTagGFP2. Bars, 18 µm. **B,** Representative images of morphological changes in NMuMG Irr- and mNT-HA-expressing cells pre and post TGF stimulation. Bars, 200 µm. **C,** Immunoblots of the indicated proteins in Irr- and mNT-HA-expressing NMuMG cells during EMT. A symbol of 15’ indicates 15 minutes. **D,** OXPHOS capacity (left) and cellular ATP levels (right) of Irr- and mNT-HA-expressing NMuMG cells during EMT. **E,** The ratio of mitochondrial DNA (mitoDNA) per nuclear DNA (NucDNA), measured by real-time qPCR. **F,** Immunoblotting of the pull-down samples for the indicated proteins. (S) and (L) indicate short and long exposure durations, respectively. **G,** Rate of wound closure by the indicated cell lines (left). Solid lines indicate simple linear regressions of these rates. Broken lines are 95% confidence intervals of the individual regression lines. The proportion of the area occupied by the cells migrating across transwells (right). (D, E, and F: Šidák’s multiple comparison test, \*\*\*\**adj. P* < 0.0001, *** *adj. P* < 0.005, ** *adj. P* < 0.01, and * *adj. P* < 0.05.)

## Discussion

It has been reported that mitoNEET is located in the outer membrane of mitochondria (2). However, some studies have suggested other possibilities, such as that mitoNEET exists in the IMM, or that a propotion of mitoNEET has a different topology towards the cytosol (20). In this study, we confirmed the location of mitoNEET using electron microscopy. Combined with the result of our experiment using MicroID2, it is indicated that mitoNEET in the OMM faces both sides (the cytosol and the inner membrane space), although the proportion of mitoNEET facing each side was not estimated. In the present study, we did not confirm whether the topological change of mitoNEET occurs during EMT, nor elucidate the mechanism by which mitoNEET is oriented towards both sides of the OMM. Nevertheless, our results suggest that the topology of mitochondrial membrane proteins has a crucial effect on biological processes, including OXPHOS, as demonstrated in the present study.

EMT confers different characteristics on cells not only in physiological conditions but also in pathological conditions such as cancer malignancy and fibrosis. Cancerous EMT has been studied extensively, and the molecular processes involved in the gene regulation and cytoskeletal rearrangements that occur have been elucidated. Analyses focusing on the metabolic background of cancerous EMT reported that ZEB1, an EMT TF, transcriptionally activates the *GLUT3* gene and promotes tumour progression by increasing glucose influx (21). Cancerous EMT has also been reported to correlate with the downregulation of OXPHOS (22). However, there are fewer studies on normal or normal-like EMT. Recently, Bhattacharya et al. showed that neural crest migration, in which EMT plays a crucial role, requires a high rate of glycolysis compared with OXPHOS (23). Our data including that from metabolomic analysis led to an alternative interpretation that EMT enhances OXPHOS to meet the ATP demands of cell motility, but because NMuMG cells are not epithelial stem cells like neural crest cells, this difference in cell origin may have caused such discrepancies. Therefore, our data from normal-like epithelial cells may provide new information for understanding non-cancer EMT.

## Contributions

H.H., Y.O., T.O., and H.S. contributed to the design of the study. H.H., S.T., K.U., T.Y., M.Y., M.W., Y.F., A.H., and S.H. performed the experiments. H.H., T.O., and H.S. analysed the data. H.H., M.M., and H.S. interpreted the data. H.H. wrote the manuscript with the contribution of T.O. and H.S.

## Acknowledgements

This work was supported by JSPS KAKENHI Grant (19K23877 and 22K15375), the Akiyama life science foundation Research Grant to H.H., and the Next-Generation Researchers Grant from Hokkaido University. This work was also supported by JSPS KAKENHI Grant Number JP22H04926, Grant-in-Aid for Transformative Research Areas ― Platforms for Advanced Technologies and Research Resources ‘Advanced Bioimaging Support’. We thank Ayae O. and Rie T. for their assistance, and H.A. Popiel for her critical reading of the manuscript.

---

**Supplementary Figure 1.**
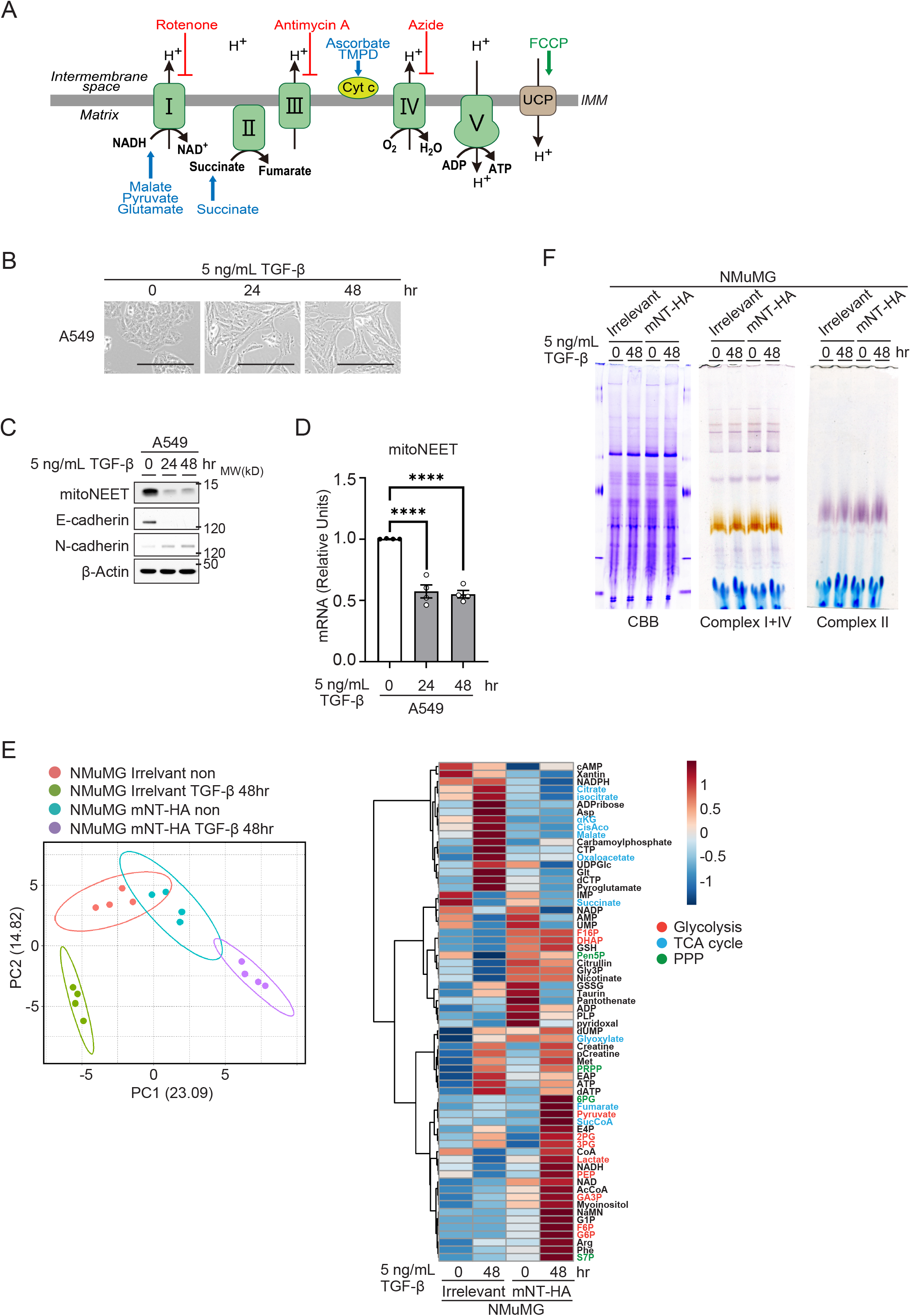
**A**, Diagram of the mitochondrial electron transfer system. The SUIT (substrate-uncoupler-inhibitor-titration) protocol used in this study is shown. Blue: substrates; red: inhibitors; and brown: uncoupler (UCP). **B,** Representative images of morphological changes in A549 cells stimulated by 5 ng/mL TGF-β. Bars, 200 µm. **C,** Immunoblots of A549 cells pre and post TGF-β stimulation. **D,** Levels of mitoNEET mRNA in TGF-β stimulated A549 cells. (C and D: Dunnett’s multiple comparison test, * *adj. P* < 0.05.) **E,** Principal component analysis of the four analysed groups (left). Heatmap of log2 fold changes of the metabolites in the four groups (right). Main metabolites of the metabolic pathways were annotated into the following three groups: glycolysis, TCA cycle, and pentose phosphate pathway (PPP). **F,** Representative images of native-PAGE analysis of isolated mitochondria from Irr- and mNT-HA-expressing NMuMG cells. Complex I + IV and complex II were detected by the in-gel assay.

